# Developmental wave of programmed ganglion cell death in human retinal organoids

**DOI:** 10.1101/2025.07.25.666895

**Authors:** Tara Brooks, Yuna K. Park, Anne Vielle, Michael Ha, Katia Del Rio-Tsonis, Michael L. Robinson, M. Natalia Vergara

## Abstract

The delicate and complex structure of the neural retina that enables proper visual function is achieved during embryonic development through a precise balance of proliferation, differentiation, and cell death. Retinal ganglion cells (RGC), the only output neurons of the retina, show a steady increase in numbers during development except for two conserved waves of developmental cell death. However, the mechanisms responsible for these phenomena in the human retina are incompletely understood. In this work, we took advantage of human induced pluripotent stem cell (hiPSC)-derived retinal organoids to explore these questions. Using different markers and quantitative techniques in three different hiPSC lines, we found a consistent decrease in RGC numbers at week 8 of differentiation, corresponding to the timing of the early developmental wave described in other vertebrates. This decrease coincided with a peak in caspase 3 activation and TUNEL(+) staining, suggesting an apoptotic mechanism. Notably, this occurred without caspase 9 activation or an increased BAX/BCL2 ratio, but with elevated caspase 8 activation, indicating involvement of the extrinsic apoptotic pathway. Together, these results show for the first time the intrinsic ability of the human retina to regulate RGC numbers through programmed cell death, providing insight into conserved developmental mechanisms and informing the use of retinal organoids in basic and translational research.

## INTRODUCTION

The retina is a complex central nervous system structure with a histoarchitecture that is fine-tuned for visual function. This cellular and structural specificity arises during embryonic development through a delicate balance between cell proliferation, differentiation and death, and a disruption in this balance can lead to congenital retinal abnormalities, and ultimately, visual impairment. Unfortunately, the mechanisms that regulate these processes to specify the correct numbers of each retinal cell type are poorly understood, particularly in the human retina.

Two waves of retinal developmental cell death have been described in animal models including zebrafish, chick, and mouse ^1-6^. The main cell type affected by these waves of cell death are retinal ganglion cells (RGC), the first cell type to differentiate in the retina and the only to relay visual information from the retina to the brain ^2,5-7^. The second wave of cell death has been more extensively studied and is hypothesized to be responsible for the refinement of the visual pathway ^2,6,8,9^. The leading hypothesis is that during RGC maturation, as these cells extend their axons to the visual centers in the brain, those who fail to make proper connections undergo programmed cell death due to the lack of retrograde neurotrophic support ^2,6,8,9^.

Conversely, the first wave of cell death is less well understood yet highly evolutionarily conserved. Studies in zebrafish, and chicks have identified apoptotic cell death as a primary mechanism responsible for this phenomenon ^3,4,6,10^, whereas recent studies in mice have suggested that this process is driven by complement-induced phagoptosis by microglia ^1^. However, while animal models have provided important insights, the mechanisms underlying RGC death and the degree of conservation of these mechanisms in humans remain unknown due to experimental limitations.

In this context, human stem cell-derived retinal organoids offer new opportunities to investigate these developmental processes. Not only do these three-dimensional retinal tissues mimic the *in vivo* retinal histoarchitecture, cellular composition and some degree of functionality, but, importantly, they have been shown to recapitulate all of the main features of retinal development following a species-specific spatio-temporal sequence ^11-17^.

In this study we took advantage of human induced pluripotent stem cell (hiPSC)-derived retinal organoids as a model to study the mechanisms of RGC death during human development. Immunohistochemical and live fluorescent reporter quantification time course studies identified a trough in RGC numbers at 8 weeks of differentiation. This developmental stage is consistent with the early wave of RGC death identified in other species, suggesting that this mechanism may be conserved in humans. This phenomenon could not be attributed to a microglia-mediated mechanism, as this cell type was absent in retinal organoids. Conversely, we found a strong correlation of caspase 3-mediated programmed cell death. Moreover, an increase in caspase 8 activation without increases in cleaved caspase 9 or BAX/BCL2 ratios suggests the involvement of the extrinsic apoptotic pathway in this process.

Thus, this study is the first to describe the intrinsic ability of the human retina to regulate RGC numbers through developmental cell death mechanisms. This has implications for the understanding of conditions that result from the dysregulation of retinal development, as well as for the application of stem cell-derived retinal organoids to disease modeling and therapeutic development.

## RESULTS

### Developmental trough in human retinal ganglion cell numbers is conserved in human retinal organoids

We used stem cell-derived human retinal organoids to investigate whether the early wave of RGC death occurs in humans. Retinal organoids recapitulate retinal structure and function, but importantly, also mimic the steps of development as they happen *in vivo* ^11-17^. Thus, they constitute an ideal model for studying retinal developmental mechanisms that affect the human retina in an *in vitro* human 3D tissue context.

We focused our analysis on a window of differentiation from weeks 6 to 10, encompassing the onset of RGC differentiation through subsequent maturation. This time frame captures the initial emergence and expansion of RGCs, beginning around day 42 of differentiation. This stage has been shown to align with early RGC specification in both human retina and retinal organoids ^11,12,15,17,18^.

To evaluate changes in the RGC population over time, we performed immunohistochemistry using HuC/D, a well-established marker of early postmitotic neurons, including RGCs ^19^. We quantified the HuC/D-positive area relative to total retinal area to measure RGC abundance across developmental time points. As expected, our results showed a steady increase in RGC numbers from week 6 to 7 of differentiation (Fig. 1A-B,F). Interestingly, at 8 weeks we observed a sudden dip in RGC numbers, followed by a steady increase in the subsequent weeks (Fig. 1C-E,F). The time of this dip in RGC numbers is suggestive of an increase in cell death and is consistent with the developmental timing of the early wave of RGC death described in other vertebrates ^1,3,4,6,10,12^.

**Figure 1.**
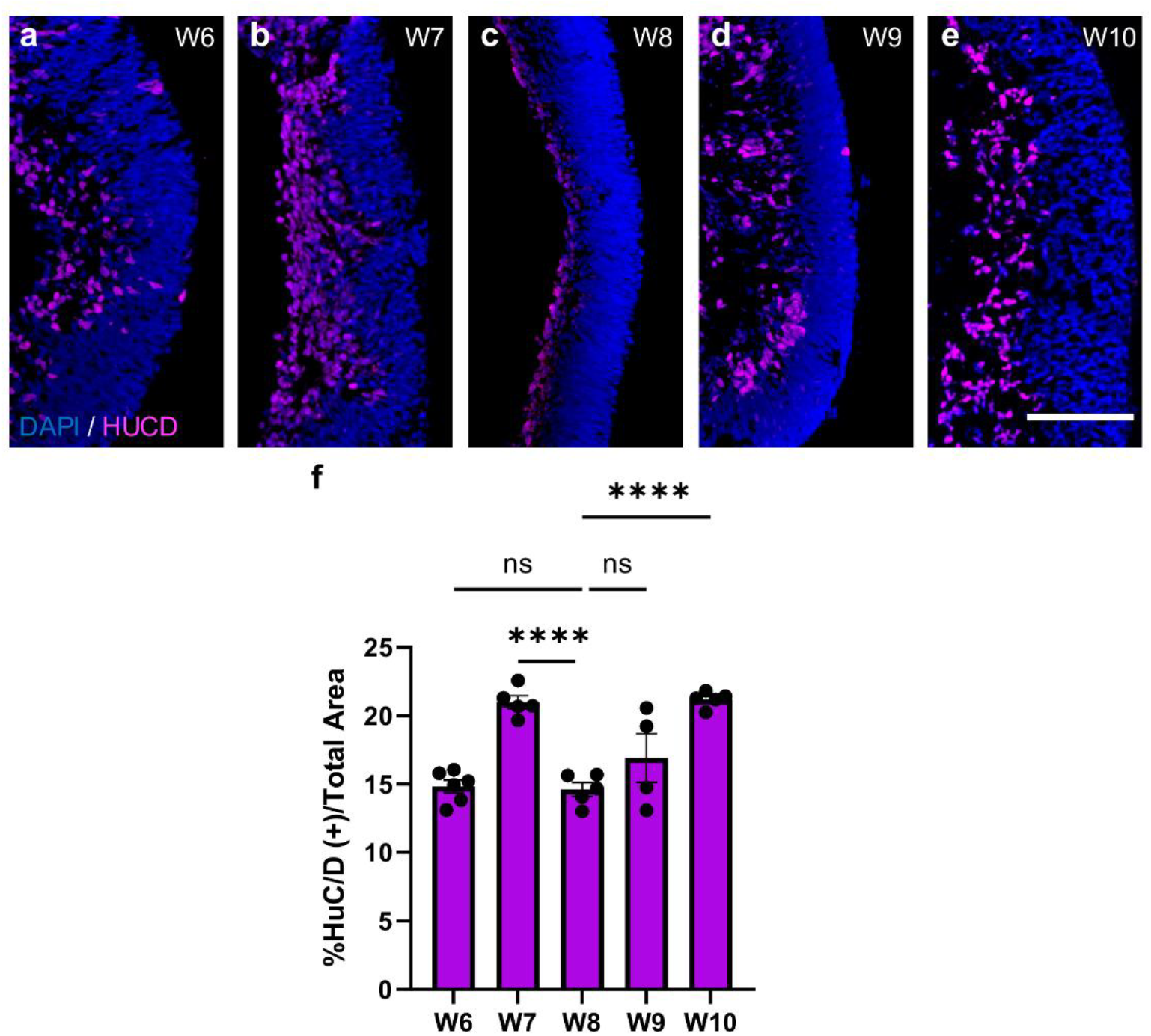
Retinal ganglion cell quantification reveals a developmental trough in human retinal organoids. a-e) Representative confocal images of retinal organoid cross sections at 20x taken on the five different differentiation time points. Sections were immunostained for the RGC marker HuC/D (red) and counterstained with DAPI (blue). f) Percentage of HuC/D(+) area/ total retinal area measured by DAPI staining. Bar graph represents mean ± SEM. N=5 biological replicates per time point as plotted. ****p<0.0001. Scale bar: 100 μm.

To validate this observation, we used the PGP1 triple-transgenic hiPSC line, which expresses eGFP under the *POU4F2* promoter; an early and specific marker of RGCs, alongside *VSX2* promoter-driven cerulean and *RCVRN* promoter-driven mCherry, as reporters for retinal progenitors and photoreceptor precursors, respectively. This enabled longitudinal tracking of specific retinal cell types in live organoids. Using the validated 3D Automated Reporter Quantification (3D-ARQ) method ^18^, we quantified fluorescent reporter expression from weeks 7 to 10. Consistent with HuC/D-based analysis, we observed a significant reduction in POU4F2-eGFP signal at week 8 (Suppl. Fig. 1), followed by recovery. In contrast, mCherry signal increased steadily during this period, suggesting the dip in eGFP expression was specific to RGCs. Moreover, this and the observed recovery in RGC numbers after week 8, suggest that the RGC decline is not due to organoid stress, necrosis or toxicity.

### Microglia are not involved in the observed trough in retinal ganglion cell numbers in human retinal organoids

In mice, microglia-mediated phagoptosis has been identified as a key mechanism underlying the early elimination of RGCs during development in mouse models ^1^. However, it has been previously reported that microglia are not present in retinal organoids, since organoid generation favors cells of neuroectodermal lineage ^11,15^. To confirm this, we examined the presence of microglia at week 8 of differentiation.

We performed immunohistochemical staining for IBA1, a well-established marker of microglia in both adult and developing human retina Retinal organoids and adult human retinal sections from cadaver eyes were processed in parallel. As expected, IBA1-positive microglia were readily detected in the adult human retina, displaying characteristic morphology (Fig. 2A–B). In contrast, no IBA1 signal was observed in any of the retinal organoids at this stage (Fig. 2C–D). Furthermore, we performed RT-PCR for CD11B and CX3CR1, two additional microglial markers. Consistent with our immunostaining results, both markers were expressed in adult human retina but undetectable in retinal organoids (Fig. 2E). These findings, which align with previous reports, indicate that the reduction in RGC numbers during early organoid development occurs independently of microglia-mediated phagoptosis.

**Figure 2.**
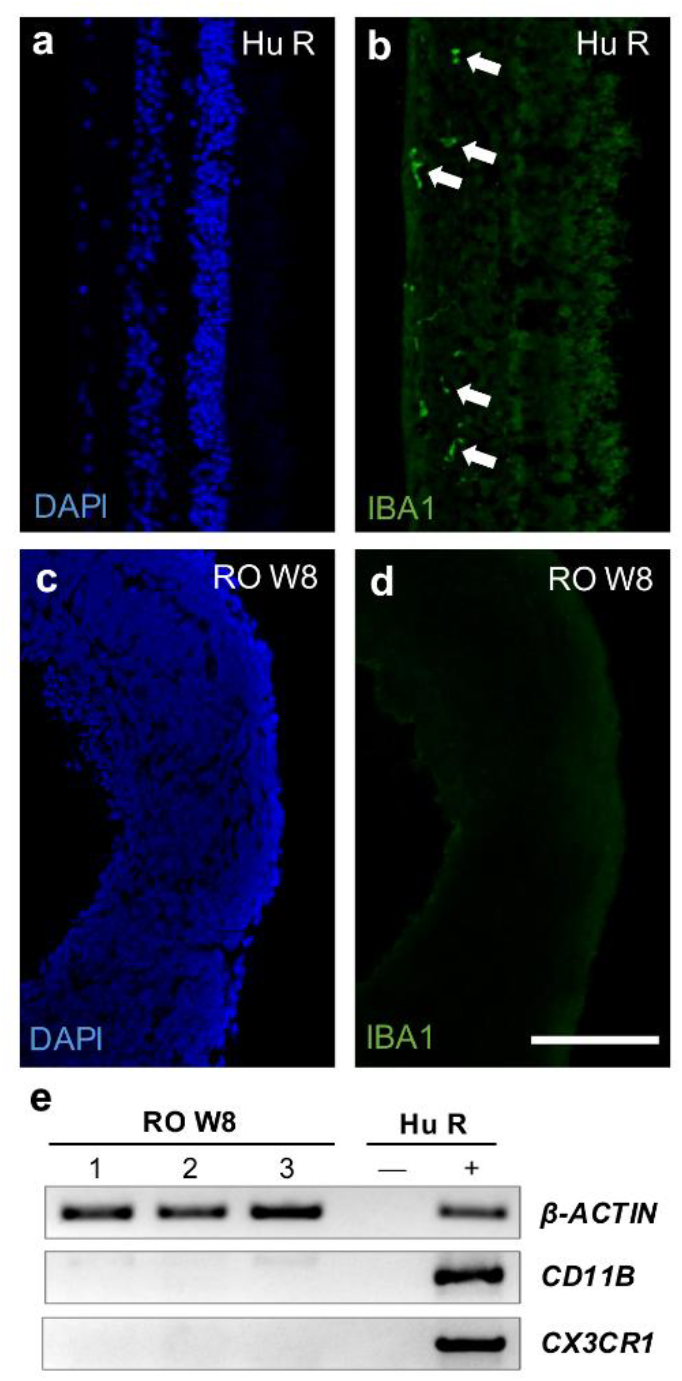
Microglia are not present in human retinal organoids. a-b) Representative confocal images of adult human post-mortem retina at 20x: DAPI (blue), IBA1 immunostaining for microglia (green). White arrows indicate positively stained microglia. c-d) Representative confocal images of retinal organoids at 10x: DAPI (blue), IBA1 immunostaining (green). e) PCR for microglia markers (CD11B and CX3CR1) on three independent human retinal organoid samples and a human retina sample (including RT(-) control) shows the absence of microglia in human retinal organoids. Scale bar: 100 μm.

### Caspase 3-dependent programmed cell death contributes to peak of developmental RGC death

The observed decrease in RGC numbers at 8 weeks of differentiation could also be explained by an increase in programmed cell death. Programmed cell death is known to be important for central nervous system development and has been found to be a major mechanism for the elimination of RGCs in other species, including zebrafish, Xenopus and chick ^3,4,6,10,20^. Additionally, caspase-mediated programmed cell death contributes to the elimination of RGCs in the adult human retina after injury and in degenerative eye diseases ^2,21^. However, whether this is the case in human retinal development has not been determined.

To study the potential involvement of programmed cell death in this phenomenon, we performed immunohistochemistry for cleaved caspase 3 on retinal organoids collected between weeks 6 and 10 of differentiation. Cleavage activates caspase 3, a major effector of both the intrinsic and extrinsic pathways of the apoptotic cascade. Our results indicate very minimal levels of caspase 3 activation between 6 and 7 weeks of retinal organoid development, followed by a sudden peak of caspase 3 activation at 8 weeks, which coincides with the sudden decrease in RGC numbers (Fig. 3A-C,F). After 8 weeks, caspase 3 activation decreases suddenly (Fig 3D-F). Notably, the majority of cleaved caspase 3 staining co-localized with HuC/D staining (Suppl. Fig. 2), suggesting that RGCs are the major cell type affected by this mechanism. To validate these results, we performed TUNEL staining in retinal organoids at similar differentiation stages. Our quantification analysis showed a similar pattern of cell death, with a peak of TUNEL(+) staining at 8 weeks of differentiation (Fig. 3G). These results support the hypothesis that caspase 3 dependent cell death plays a major role in the trough of RGC numbers observed at this time point.

**Figure 3.**
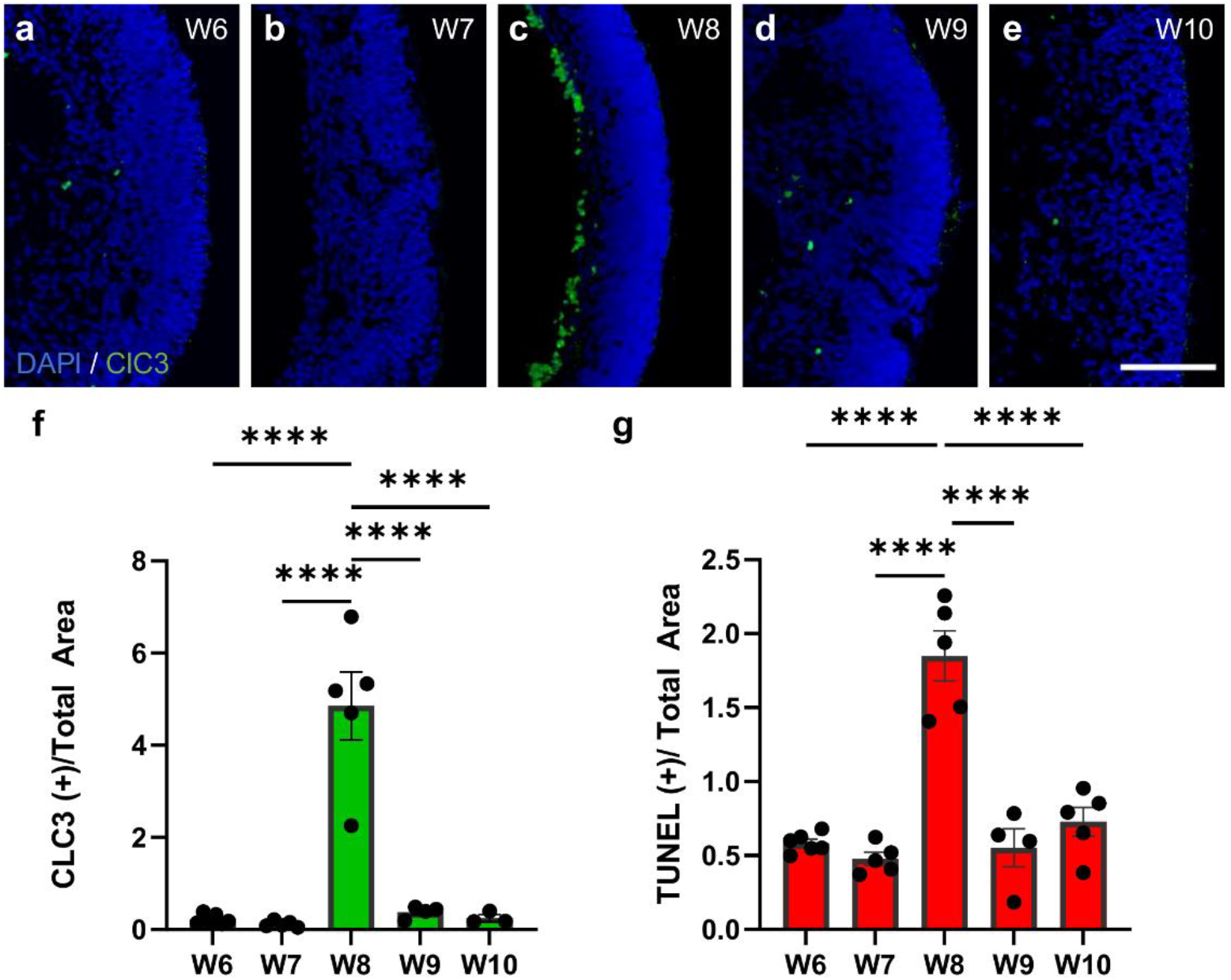
Caspase-mediated programmed cell death is highest at the time of RGC loss. a-e) Representative confocal images of retinal organoid cross sections at 20x taken on the five different collection time points post-differentiation: DAPI (blue), cleaved caspase3 (ClC3; green). f, Percentage of ClC3(+) area/ total area measured by DAPI at each collection time point. g) Percentage of TUNEL(+) area/ total area measured by DAPI at each collection time point. N=5 biological replicates per time point as plotted. Bar graph represents mean ± SEM. ****p<0.0001. Scale bar: 100 μm.

### Involvement of the extrinsic apoptotic pathway in RGC death in retinal organoids

To further investigate the mechanisms underlying programmed cell death during early retinal development, we examined the expression of key apoptotic markers in retinal organoids. Western blot analysis was performed on retinal organoids at week 8 of differentiation—when RGC loss is most pronounced—and compared to week 7, when cell death remains minimal.

Consistent with our immunofluorescence findings, we observed a marked increase in cleaved caspase-3 levels at week 8 (Fig. 4A), confirming active apoptosis. Surprisingly, however, the Bax/Bcl-2 ratio decreased at this stage (Fig. 4A–B), suggesting a shift toward an anti-apoptotic state within the intrinsic pathway. In line with this, we found no evidence of caspase-9 cleavage, indicating that the intrinsic apoptotic cascade is not the primary driver of cell death at this time point.

**Figure 4.**
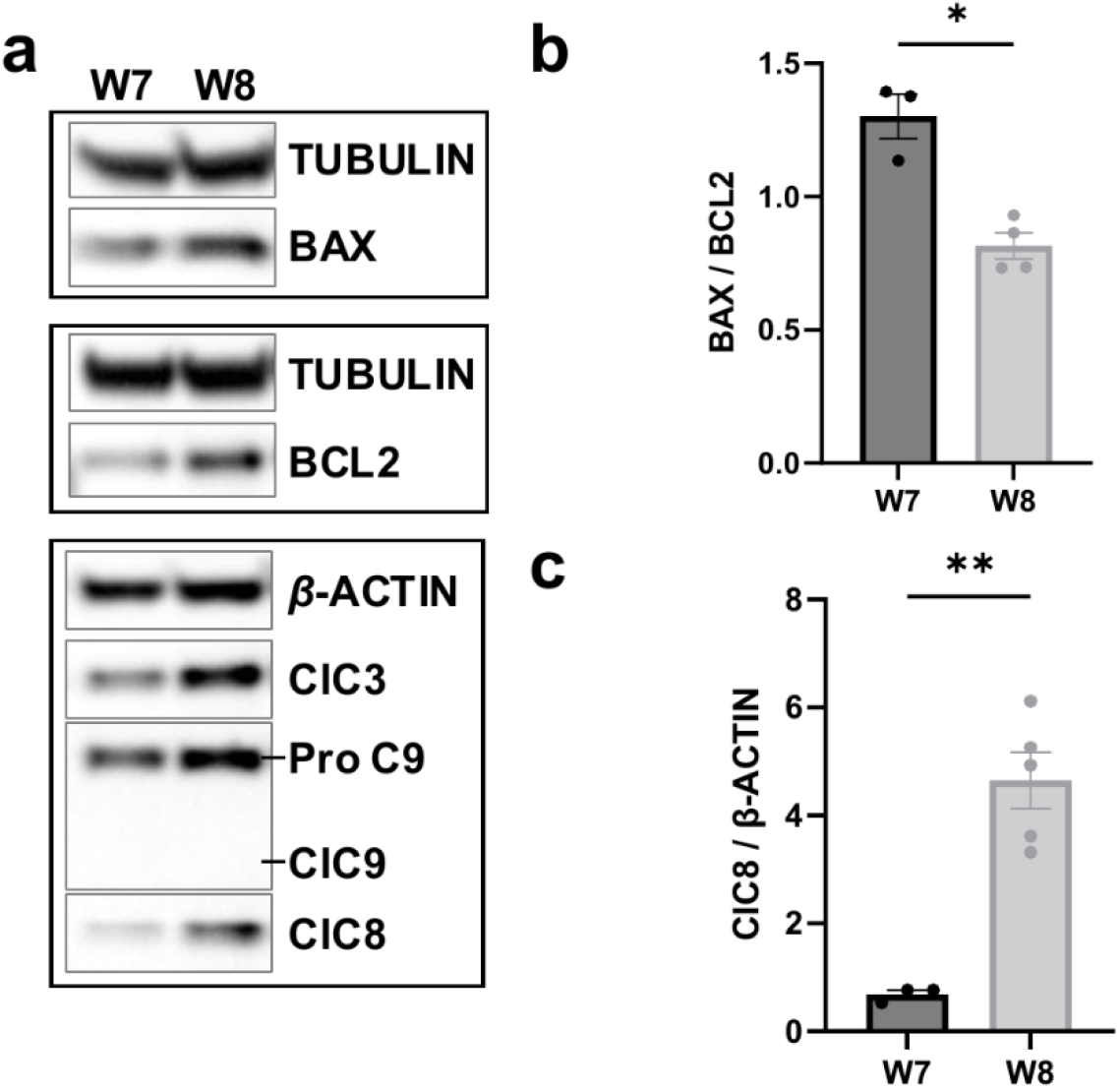
Analysis of apoptotic pathway components in retinal organoids. a) Representative Western blot images show expression of cleaved caspase-3 (ClC3), cleaved caspase-8 (ClC8), cleaved caspase-9 (ClC9), BAX, and BCL2, in retinal organoids at weeks 7 and 8 of differentiation. Notice the lack of ClC9 detection. b) Western blot quantification shows a decrease in the BAX/BCL2 ratio at 8 weeks of differentiation-the time of RGC death. c) Western blot quantification also shows an increase in ClC8 at 8 weeks of differentiation. N=3-8 biological replicates per time point, as plotted. Bar graph represents mean ± SEM. *p<0.05; **p<0.01. Scale bar: 100 μm.

Instead, we observed a significant increase in cleaved caspase-8, a hallmark of extrinsic apoptotic pathway activation (Fig. 4A, C). These findings suggest that the early wave of RGC death in human retinal organoids is mediated primarily through extrinsic apoptosis, although the specific death ligands and receptors responsible for initiating this pathway remain to be identified.

## DISCUSSION

The retina is a neural tissue whose function is dependent on the finely tuned complexity of its structure. This structure arises during development through a delicate balance of cell proliferation, differentiation, and death ^7^. Despite the crucial importance of this balance for producing a tissue with the right proportions of different cell types, relatively very little is known about how retinal cell numbers are regulated during development. In this work, we focused on how programmed cell death affects the development of RGCs in the human retina.

Two waves of cell death primarily affecting RGCs have been described in vertebrate retina development ^1-6,10,20^. These waves seem to be highly evolutionarily conserved, suggesting that they play an important developmental role. The later wave of RGC death is thought to represent a circuit refinement, as it takes place when RCGs are making their synaptic connections to their brain targets ^2,6,9^. The leading hypothesis is that when correct synapses are formed, those circuits are strengthened by retrograde neurotrophic support that promotes the survival of RGCs, and the cells that fail to make connections undergo cell death ^2,6,9^. This is represented in retinal organoid models by the depletion of retinal ganglion cells that occurs between 15 and 18 weeks of differentiation ^11,15^. Moreover, a recent study on assembloids—models made of a retinal organoid linked to a brain organoid—revealed that RGCs were able to extend their axons to the brain organoid and showed increased survival rates. This finding further suggests that this process is comparable to the second wave of RGC death observed in vertebrates ^22^.

In contrast, the earlier wave of developmental RGC death is poorly understood, although its evolutionary conservation suggests its importance as a developmental mechanism. Moreover, this phenomenon has never been investigated in humans. This gap in the knowledge prompted us to investigate the timing, pattern and mechanisms responsible for this phenomenon using human retinal organoids as a model.

Our results using organoids derived from three human iPSC lines and evaluated using different RGCs markers and different methods of quantification, showed a steady increase in RGCs from 6 to 7 weeks of differentiation, followed by a sharp drop at week 8 (day 56). Subsequently, the number of RGCs began to steadily increase from weeks 9 to 10 of differentiation. The timing of this trough in RGC numbers is developmentally equivalent to that of the first wave of RGC death in other species ^1,3,4,6,10,12^, suggesting the conservation of the first wave of RGC death in human retinal organoids.

When exploring the mechanisms responsible for this phenomenon, our results did not support a microglia-mediated mechanism, which had been reported in mice ^1^. Specifically, we did not find evidence of microglia presence in our retinal organoids, consistent with prior reports ^11,15^. Conversely, the sharp increase in RGC loss was highly correlated with a peak in caspase 3 activation and TUNEL-positive signal, pointing to caspase 3-dependent programmed cell death as a mechanism for RGC elimination in human retinal organoids. Moreover, activation of caspase 8 with concomitant lack of caspase 9 activation and a low Bax/Bcl2 ratio, suggested the involvement of the extrinsic apoptotic pathway. However, the ligands that activate this pathway in retina development remain to be elucidated. From these experiments we cannot discard a possible microglial involvement in this phenomenon *in vivo*, which is a limitation of this study. However, the fact that this wave of RGC death occurs consistently in human retinal organoids indicates that programmed cell death is endogenous to the developing retinal tissue, and thus likely to occur *in vivo* as well. The discrepancies with previous studies could be attributed to species differences. It is also possible that programmed cell death co-exists with microglia-driven phagocytosis in the human retina, though this cannot be confirmed with the current model.

In summary, our work sheds light onto the processes that shape the human retina during development, something that has been poorly explored in the past due to a lack of adequate models, and further supports the usefulness of human retinal organoids to study human retinal development. This research has important implications for our understanding of conditions that result from the dysregulation of retinal development, such as what occurs in Down syndrome, a condition driven by the triplication of chromosome 21 that presents with a supernumerary retina ^23^. Finally, these results may also have implications for the application of retinal organoid models in regenerative therapy and pharmacological testing.

## METHODS

### Human induced pluripotent stem cell culture

Human iPSC lines used in this study were the cord blood-derived episomal hiPSC line A18945 (Gibco), the RNA-reprogrammed, urine-derived hiPSC line iLC67-3 generated at the University of Colorado, and the triple transgenic reporter hiPSC line PGP1 generated by CRISPR/Cas9-mediated homology-directed repair (HDR) to replace stop codons of the VSX2, POU4F2, and RCVRN genes P2A sequences fused to Cerulean, green fluorescent protein, and mCherry reporter genes respectively ^24^. All iPSC lines were obtained commercially or through a material transfer agreement in a de-identified fashion. This study does not qualify as human subject research. Providers guarantee that lines were generated under IRB approval at the corresponding institutions. Quality control of cell lines included karyotypic analysis and routine mycoplasma testing by PCR. hiPSCs were maintained on Matrigel (growth-factor-reduced; BD Biosciences)-coated plates with mTeSR1 medium (Stemcell Technologies) according to WiCell protocols. Cells were passaged at ∼70% confluence. Colonies containing differentiated cells were marked and mechanically removed before passaging. The use of hiPSCs in this study conforms to the University of Colorado Institutional Biosafety Committee (IBC).

### Retinal organoid differentiation

Stem cells were maintained and differentiated into retinal organoids as previously described ^17^. Briefly, hiPSCs were cultured on mTeSR media following manufacturer’s specifications. On day 0 of the differentiation protocol, hiPSC colonies were lifted using dispase (Sigma), broken into small clumps by gentle pipetting, and cultured in suspension in a 37°C cell culture incubator with 5% CO_2_ to form embryoid bodies (EBs). Over the next 3 days they were gradually transitioned to neural induction medium containing DMEM/F12 (1:1), 1% N2 supplement (Invitrogen), 1 x minimum essential media-non essential amino acids (NEAAs), 2ug/ml heparin (Sigma). On day 7 the EBs were seeded on Matrigel coated plates. On day 16 they were transitioned to retinal differentiation medium (DMEM/F12 (3:1) supplemented with 2% B27 (without vitamin A, Invitrogen), 1x NEAA and 1% antibiotic–antimycotic (Gibco)). Between days 20-24 retinal domains were identified using phase contrast microscopy and mechanically and manually lifted from their plates using tungsten needles. The lifted retinal domains were grown in suspension and allowed to fold into retinal cups. Media was switched to DMEM/F12 (3:1) supplemented with 2% B27, 1x NEAA, and 1% antibiotic–antimycotic on day 30. On day 50, 1µM retinoic acid was added to the media daily.

### Tissue collection and processing

Retinal organoids were collected weekly between weeks 6 and 10 of differentiation (42, 49, 56, 63 and 70 days of differentiation). Organoids were washed in phosphate buffer solution (PBS) and fixed in 4% paraformaldehyde for 10 minutes. After fixation, organoids were transitioned through a sequential overnight incubation in 6.75%, 12.5% and 25% sucrose in 1x PBS, in order to equilibrate the osmotic pressure, before being embedded in Sakura Tissue-Tek OCT compound. The embedded organoids were frozen and stored at -80 °C until sectioned. Sections were taken on a Microm cryostat at -24 °C, at 12 µm in thickness. A de-identified human eye from a cadaveric donor (female, 65 years old) was obtained from the Miracles in Sight Eye Bank (North Carolina), fixed in 10% (w/v) methanol-free neutral buffered formalin for 2 hours, cryoprotected in a sucrose gradient and embedded in Sakura Tissue-Tek OCT compound for cryosectioning at 12 µm thickness. All procedures involving human tissue were conducted in accordance with the Declaration of Helsinki and were deemed exempt due to use of de-identified cadaveric tissue. For protein analysis, retinal organoids were rinsed in PBS and flash-frozen in liquid nitrogen. Tissues were then lysed in RIPA buffer (Sigma) with protease inhibitors using a homogenizer, and centrifuged for 15 minutes at 14,000g. Protein concentration in the supernatant was determined using the Biorad DC Protein assay (BioRad).

### Immunofluorescence

Three slides per retinal organoid were immunostained for each marker analyzed. For microglia analysis, human post-mortem retinal slides were processed alongside differentiation week 8 (day 56) retinal organoids. Slides were washed in PBS three times for five minutes, then blocked for 1 hour in 10% normal donkey serum diluted in PBS 0.25% Triton X-100 (PBST) at room temperature (RT). Slides were then incubated overnight at 4°C in primary antibody solution diluted in 2% normal donkey serum with 0.25% PBST. Primary antibodies used were mouse-anti-HuC/D antibody at 1/100 (Invitrogen, catalogue number A21271), rabbit-anti-IBA1 at 1/100 (Proteintech, cat# 10904-1-AP, AB_2224377) and rabbit-anti-cleaved caspase 3 antibody at 1/100 (Cell Signaling, cat# 9664, AB_2070042). Slides were then washed three times in PBS for five minutes and incubated for 1 hour at RT in the corresponding secondary antibodies diluted 1/1000 in 2% normal donkey serum with 0.25% PBST. Secondary antibodies used were donkey-anti-mouse Alexafluor 546 (Invitrogen, cat# A10036) and donkey-anti-rabbit Alexafluor 488 (Invitrogen, cat# A-21206). Slides were washed for five minutes in PBST, followed by two ten-minute washes in PBS, and mounted in DAPI mounting medium (SouthernBiotech DAPI Fluromount, cat# 0100-20) with a coverslip.

### Imaging and image analysis

Images were taken on a Nikon C2 confocal microscope and stitched together using Nikon Element software. Images were analyzed using Fiji/ImageJ. Five organoids were analyzed per collection time point for each marker (biological n=5). For each organoid, three slides from the middle of the organoid were imaged and analyzed (technical n=3), to minimize technical error. For each fluorescence microscopy image, a rectangular region of dimensions 180um x 220um was selected. In order to approximate RGC numbers, we measured the area of HuC/D positive immunostaining and expressed it as a ratio to the area of DAPI positive staining in the same region. To approximate the amount of programmed cell death we performed a similar analysis using the area of cleaved caspase 3-positive immunostaining to DAPI staining. Ratios were consistent across the three images per organoid per collection time point. The ratios from each slide were converted into percentages and the percentages from the three slides from each organoid were averaged.

### 3D Fluorescent reporter quantification

Longitudinal quantification of fluorescent reporters in live organoids was assessed using 3D-Automated Reporter Quantification as previously described ^18,25^. Briefly, organoids derived from the transgenic PGP1 hiPSC line were plated in clear u-bottom 96 well plates at 30 days of differentiation and cultured in these conditions thereafter. Total fluorescence intensity was assessed weekly between 7 and 10 weeks of differentiation using a Tecan Spark plate reader. Before each reading, organoids were washed twice in PBS and changed to DMEM FluoroBrite (Thermo Fisher Scientific). The instrument was pre-warmed to 37°C and top-reading mode was selected. After z-dimensional focusing, fluorescence intensity was read in 3×3 fields at the center of each well using 20 flashes and the following excitation and emission parameters: Cerulean (ex: 435nm/ em: 478nm), mCherry (ex: 567nm/ em: 625nm), eGFP (ex: 476nm/ em: 516nm), with a bandwidth of 10 nm. Wells with media alone were used for background subtraction and wild type retinal organoids of the same age were used as negative controls. Cerulean expression was as an approximation for size normalization within each time point.

### Protein quantification

Protein levels were assessed by Western blot. Eight organoids were pooled per sample and 4 independent biological samples were analyzed per time point. Organoids were lysed in RIPA buffer (Sigma Aldrich) containing protease inhibitors (Roche). Protein concentrations were determined using the DC Protein Assay (BioRad). Proteins were separated in a 12% SDS-PAGE gel and transferred to PVDF membranes using the iBlot2-System (Life Science Technologies). Membranes were blocked in 1X casein for 1 hour followed by incubation with primary antibodies overnight at 4°C as indicated in Supplementary Table 1. Blots were then washed and incubated with HRP-conjugated secondary antibodies (supplementary table 1) for 30 minutes at room temperature. SuperSignal West Femto kit (Thermo Scientific) was used for chemoluminescent detection.

### RT-PCR

Total RNA was purified from retinal organoids collected at week 8 using the RNAqueous Micro Kit (Invitrogen, cat#AM1931). Total RNA was also purified from the retina from a 52-year-old human female using the Direct-zol RNA Mini Prep (Zymo Research, cat#R2050S). Reverse transcription was performed using the Maxima H Minus cDNA Synthesis Master Mix with dsDNase (Thermo Scientific, cat#M1669). One microgram of cDNA from each sample was mixed with 0.125µL of BioReady rTaq, 0.25µL dNTP mix, 1.25µL of 10X reaction buffer, 9.375µL of water, and 0.5µL of a pooled mixture of either CD11B primer mix (forward: 5’-ATG GCT CTC AGA GTC CTT CT –3’; reverse: 5’-CTG CAT CAA AGA GAA CAA GGT TT-3’, 10µM each), CX3CR1 primer mix (forward: 5’-GAA GAG CTC TCT GGC TTC TG-3’; reverse: 5’-TTC AGG CAA CAA TGG CTA AAT G-3’, 10µM each), or Beta-actin primer mix. Thermocycler settings were: 1 cycle of 94°C for 5 minutes, 35 cycles of 94°C for 30 sec, followed by either 54°C (for CD11B) or 61°C (for CX3CR1) for 30 seconds, 1 cycle of 72°C for 1 minute, and 1 cycle of 72°C for 5 minutes. No-RT and non-template controls were included. Amplified products were resolved a 2% agarose gel and visualized using ethidium bromide staining.

### Statistical Analysis

Statistical analysis was performed using one-way ANOVA with Post Hoc Tukey HSD test using Statpages (https://statpages.info/anova1sm.html) and Jamovi (https://www.jamovi.org/). A normal distribution of the data was determined for all experiments. Confidence interval of 95% was selected for significance purposes. A P-value of <0.05 was considered significant. Bar graphs represent the mean +/-SEM.

## AUTHOR CONTRIBUTIONS

MNV conceived and designed the study. TB, YKP and AV performed the experiments. TB, YKP, AV, MH and MNV analyzed the data and performed statistical analyses. MNV, TB and YKP interpreted the data. KDRT and MLR developed the PGP1 triple fluorescent reporter line. MNV and TB wrote the manuscript. YKP, MH and MNV edited the manuscript. All authors contributed to manuscript revision, read, and approved the submitted version.

## DATA AVAILABILITY

All data generated or analyzed during this study are included in this published article and its Supplementary Information files. Additional raw datasets supporting the findings of this study are available from the corresponding author upon reasonable request.

