## Supplementary figures and tables for "Developmental wave of programmed ganglion cell death in human retinal organoids"

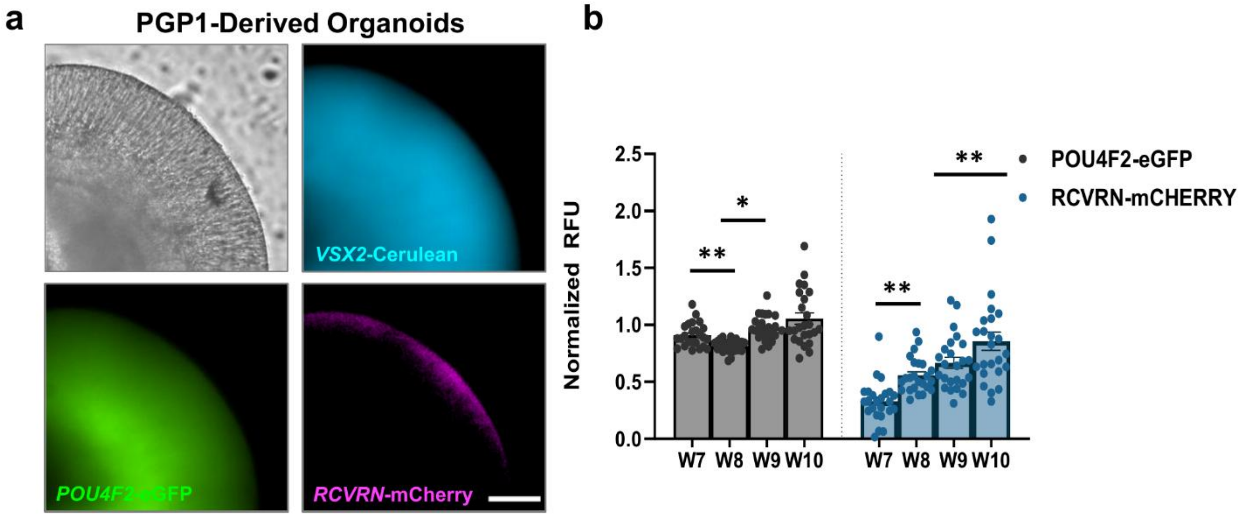


**Figure S1. Longitudinal quantification of fluorescent reporters in live retinal organoids.** a) Fluorescence micrographs of whole-mount live transgenic organoids at 10 weeks of differentiation show expression of *VSX2*-Cerulean (retinal progenitors), *POU4F2*-eGFP (RGCs), and *RCVRN*-mCherry (photoreceptor precursors). b) Quantification of reporter expression normalized with VSX2-Cerulean. Bar graph represents mean ± SEM; individual samples are plotted. *p<0.05; **p<0.01. Scale bar: 100 μm.

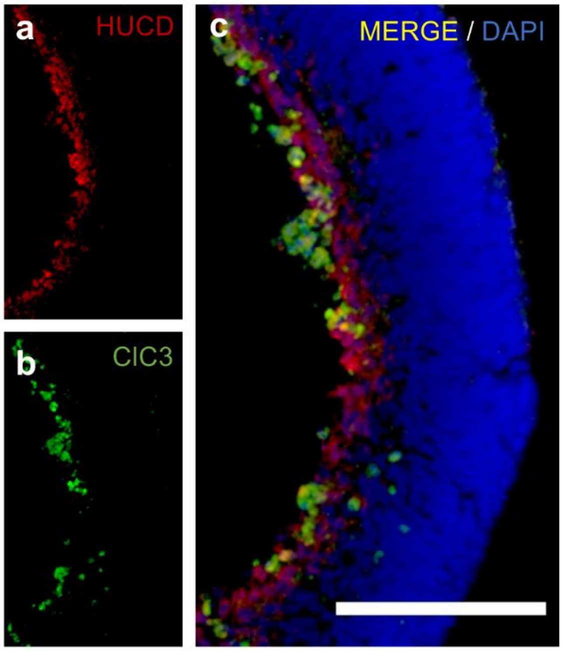


**Figure S2. Developmental wave of cell death primarily affects RGCs.**
Fluorescence micrographs of RO cryosections at 8 weeks of differentiation. a) Immunofluorescent staining for HuC/D (red) labels RGCs. b) Immunofluorescent staining for cleaved caspase3 (ClC3; green) labels cells undergoing apoptosis. c) Merged image showing co-localization of these markers. Scale bar: 100 µm.

**Supplementary table 1. Antibodies for Western blot analysis**

| Antibody | Source | Dilution |
|---|---|---|
| E7 (Anti-β Tubulin) | Developmental Studies Hybridoma Bank | 1:3000 |
| β-actin (C4) | Santa Cruz Biotechnology (cat# sc-47778) | 1:5000 |
| Anti-BAX | Proteintech (cat# 50599-2-Ig) | 1:12000 |
| Anti-BCL2 | Proteintech (cat# 60178-1-Ig) | 1:2500 |
| Anti-Caspase 3 | Cell Signaling (cat# 9662s) | 1:1500 |
| Anti-Caspase 8 | Proteintech (cat# 66093-1-Ig) | 1:5000 |
| Anti-Cleaved Caspase 9 | Cell Signaling (cat# 7237P) | 1:1000 |
| Peroxidase Labeled Anti-Rabbit IgG (H+L) | Vector (cat# PI-1000) | 1:3000 |
| Peroxidase Labeled Anti-Mouse IgG (H+L) | Vector (cat# PI-2000) | 1:3000 |
